# Long-term cytotoxic NK cells with broad anti-tumour capacity proliferate selectively, without exhaustion, after BCG priming and extremely low doses of cytokines

**DOI:** 10.1101/2023.06.07.543999

**Authors:** María-José Felgueres, Gloria Esteso, Álvaro F. García-Jiménez, Ana Dopazo, Luis Martínez-Piñeiro, Hugh T. Reyburn, Mar Valés-Gómez

**Author notes:** Corresponding author. Mar Valés-Gómez; CNB-CSIC Darwin, 3, 28049 Madrid. The authors declare no potential conflicts of interest.

## Abstract

**Background:** Natural killer (NK) cell-based immunotherapies, currently under investigation, appear to be safe, efficient treatments in patients with haematological tumours. Nevertheless, the short-lived nature of these cells combined with the need to infuse large number of cells for efficient tumour elimination represent important challenges for the development of NK cell-based therapies. Although NK cell anti-tumour activity is regulated by cytokines, constant stimulation together with the immunosuppressive tumour environment can result in NK cell exhaustion. Therefore, improved approaches to produce highly cytotoxic and longer-lived NK cells are of considerable clinical interest.

**Methods:** Peripheral blood mononuclear cells (PBMC) are primed *in vitro* with a pulse of either *Bacillus Calmette-Guérin* (BCG) vaccine or a cell wall extract of *M. bovis*, followed by weekly stimulations with low doses of IL12, 15 and 21. The phenotype and anti-tumour fitness of the activated NK cell culture were examined using scRNA-seq, flow cytometry and functional assays, including degranulation, specific cytotoxicity and IFNγ release.

**Results:** we describe a novel strategy for the generation of long-lived activated NK cells capable of killing a broad range of solid tumours. A unique subset of cytotoxic NK cells (CD56^high^ CD16^+^ NKG2A^+^) specifically proliferated *in vitro*, and was further expanded without functional exhaustion under minimal survival cytokine combinations. Mycobacterial cell-wall fractions also activated NK cells that recognised tumours efficiently, and proliferated well, and this approach has the advantage that no live bacteria are present in the cultures.

**Conclusions:** We propose that BCG-priming to expand anti-tumour NK cells, without cell sorting, could be a scalable and economical basis for the development of safe and universal cellular immunotherapies against solid tumours.

**Key messages:** Adoptive therapy with sorted NK cells grown in IL12, 15, 18 are being tested in clinical trials, but are only efficient for haematological tumours. In addition, their survival *in vivo* is limited. Here, we define culture conditions that drive the selective proliferation of long-lived natural killer (NK) cells, without the need of cell sorting, in minimal doses of cytokines, after priming with BCG or mycobacteria components. BCG-primed NK cells grow and maintain effective cytotoxic function against a variety of solid tumours *in vitro*, without exhaustion for at least 28 days of culture. This new approach provides the basis for the generation of innate adoptive cell therapy tools.

## Background

Natural killer (NK) cells are cytotoxic lymphocytes of innate immunity with potent anti-tumour capacities that make them potential efficient agents in cancer immunotherapy (1, 2, 3). Specifically, the combination of efficient cytotoxic tumour recognition, in the absence of complications like cytokine release syndrome, neurotoxicity or graft-versus-host disease (GVHD) (4) make them attractive candidates for cellular therapy of cancer. Recent trials of adoptive NK cell therapy, reported complete remissions of 17-50% of haematological cancers, but many fewer responses were noted in solid tumour trials (2). These observations have spurred the search for new protocols for *ex vivo* NK cell priming, using multiple cell sources and cytokine-based enhancement of effector functions to try and develop universal “off-the-shelf” NK cell therapies. NK cells generated with these strategies are currently being evaluated in several phase I/II clinical trials against both haematological and solid tumours.

NK cells are an extremely heterogeneous population (5, 6) and only certain subsets are active for tumour elimination. In peripheral blood, the majority of circulating human NK cells have low expression of CD56. This CD56^dim^ population, considered the mature cytotoxic NK cell subset, usually expresses high levels of CD16 (FcγRIII) and can also mediate antibody-dependent cellular cytotoxicity (ADCC) (7). CD56^bright^ NK cells, generally less cytotoxic, express little or no CD16, and are considered regulatory. CD56^dim^ and CD56^bright^ NK cells express different levels of several activating and inhibitory receptors including KIR, CD94/NKG2 heterodimers, and NCRs among others, contributing to the balance of signals that regulate NK function [for review (8)]. This heterogeneity of NK cell phenotypes which, in many instances complicates our understanding of their biology, has the advantage of providing a very versatile immune cell type that can be used for a large variety of specific functions, including the elimination of pathogen-infected and cancer cells.

Studies of NK cell education and differentiation in secondary lymphoid tissues, have revealed the strong influence exerted by cytokines, such as IL15, IL12, IL18, IL21 and IFNα/β for NK maturation and function (9, 10, 11). The use of these cytokines in *vitro*, has allowed characterization of distinct phenotypes and functions of NK cells. IL12 alone sustains NK cell viability without proliferation and acts synergistically with IL18 and IL21 to stimulate IFNγ production (12, 13, 14). IL15 is essential for both NK differentiation and survival and considerably enhances the anti-tumour response of activated NK cells (15). So, brief exposure of NK cells to high doses of the cytokines IL12, IL15 and IL18 results in the upregulation of IFNγ, perforin and granzymes (16).

All this knowledge on NK cell biology has contributed to both the improvement of NK cell culture protocols (17) and has also been translated into therapies in recent years. Cytokine-induced memory-like (CIML) NK cells produce IFNγ in response to tumour cells, upon cytokine re-stimulation (18, 19) and have been tested in phase I and phase II (20, 21, 22) clinical trials. Leukaemia patients successfully tolerated CIML cell infusions and 4/8 paediatric patients achieved complete remission. Other “adaptive-” or “memory-like” NK cells were first identified in the context of human cytomegalovirus (HCMV) infection (23, 24, 25). NKG2C^+^ NK cells are long-lived and produce IFNγ after CD16 stimulation. However, many questions about the origin and function of these NKG2C^+^ cells remain unanswered (26). A third type of memory-like activity described for innate immune cells, referred to as trained immunity, corresponds to the non-specific protection to unrelated secondary challenges elicited by vaccination with *Bacillus Calmette-Guérin* (BCG)(27). For example, NK cells isolated from healthy individuals who received BCG vaccination produced more pro-inflammatory cytokines *ex vivo* upon re-challenge with either similar or unrelated pathogens, even three months after first encounter (28). Interestingly, direct intravesical instillations of BCG is one of the first successful cancer immunotherapies used for decades for the treatment of non-muscle invasive bladder cancer (NMIBC) (29) and we have demonstrated that BCG-activated NK cells kill bladder cancer cells efficiently (30). In this BCG-priming model for bladder cancer, the activated NK cells expressed CD16, KIR, CD57 and CD94/NKG2A. This phenotype was consistent with an anti-tumour CD56^high^CD16^+^ phenotype previously described for cytokine-activated NK cells (31) and is also reminiscent of CIML-NK cells (19).

Whether anti-tumour BCG-stimulated NK cells have features of CIML, trained immunity or adaptive-like NK cells is still unknown. We demonstrated that cytokines were a crucial step for NK cell activation in the context of BCG (30), but very low levels of soluble factors were released from BCG-activated PBMC *in vitro*. However, these almost undetectable concentrations were sufficient to stimulate CD56 upregulation on NK cells and activate their cytotoxic function against tumours. In parallel, cytokines detected in urine of BCG-treated bladder cancer patients usually decrease quickly and only certain chemokines remain detectable several days after instillations (32). These findings suggest that extremely low cytokine concentrations might be sufficient for effective immune cell activation. We hypothesised that, after BCG-priming activated NK cells could grow further using minimal doses of cytokines, so that activation of immune effector cells would not have detrimental effects of excessive activation that, in many instances, can result in cell anergy. Moreover, BCG could provide beneficial priming and complement cytokine activation for anti-tumour responses, since to date, neither cytokines nor cytokine antagonists have been effective as monotherapies in patients with advanced-stage cancers, and in some cases, have even caused toxicity (33).

Here we tested whether BCG-primed anti-tumour lymphocytes were able to maintain their phenotype and function after stimulation with minimal doses of cytokine combinations, so that NK cells could grow without becoming exhausted. We show that anti-tumour BCG-primed NK cells proliferate *in vitro* after weekly addition of nearly negligible doses of IL12, 15, and 21. The simple protocol described here dramatically increased the total number of NK cells (200-fold) which, remarkably, maintained an effective anti-tumour function and killed a broad range of solid tumour cell lines, not just bladder cancer. The presence of activating receptors was enough to counteract the putative inhibitory capacity of NKG2A. Incubation with cell wall fractions of *M. bovis*, also primed NK cells that proliferated further upon cytokine culture. Thus, an innovative method, based on an FDA-approved immunotherapeutic agent together with optimised cytokine concentrations, provides a model for enhanced NK functional fitness that could allow novel regimes for cancer cell therapy.

## Methods

### BCG, cell wall fragments and peripheral blood populations

PBMCs from buffy coats of healthy donors were obtained from the Regional Transfusion Centre, Madrid, with informed consent from the participants and with the ethical permission and experimental protocols approved by local and CSIC bioethics committees. All methods were carried out in accordance with biosafety guidelines and regulations authorized by CNB-CSIC. PBMCs were isolated by centrifugation on Ficoll-HyPaque and cultured in complete (2 mM L-glutamine, 0.1 mM nonessential amino acids, 1 mM sodium pyruvate, 100 U/mL penicillin, 100 U/mL streptomycin, 10 mM Hepes, 50 µM β-mercaptoethanol) RPMI-1640 medium (Biowest) supplemented with 5% FBS (Capricorn), 5% HS (Sigma).

BCG-Tice strain (OncoTICE, MSD) (2% alive bacteria) (34) aliquots were reconstituted in RPMI 10% DMSO and stored at -80LJC. *Mycobacterium bovis*, Strain AF 2122/97 (ATCC® BAA-935™), Cell Wall Fraction NR-31213, was obtained from BEI Resources, NIAID, NIH.

### BCG-mediated stimulation

PBMC co-culture with BCG *in vitro* model was described previously (30, 34). Briefly, 10^6^ PBMC/ml were incubated in 24-well plates with or without BCG at a 6:1 ratio (total bacteria to PBMC). After one week in culture, cells in suspension were either recovered from the co-culture, centrifuged, and analysed by flow cytometry or kept in culture for another week, as indicated.

### Cell lines

All cell lines were genotyped for authentication at the genomics service of the Instituto de Investigaciones Biomédicas (IIBB-CSIC, Madrid). The bladder cancer cell lines T24, J82, and RT-112 were previously described (30). Human metastatic melanoma cell lines Ma-Mel-86c and Ma-Mel-86f, provided by Prof. Annette Paschen (University Hospital of Essen, Germany), were described previously (35, 36). The MCF7 and MDA-MB-453 breast adenocarcinoma and metastatic carcinoma, the SW480 colon adenocarcinoma, the MKN45 gastric cancer, K562 erythroleukaemia cells and H3122 lung cancer cell lines were cultured in complete RPMI-1640 medium (Biowest) supplemented with 10% FBS. All cells were kept at 37°C, with humidified atmosphere of 5% CO_2_. Cells were regularly tested for mycoplasma contamination.

### Flow cytometry

Cells were washed with PBA [PBS supplemented with 0.5% bovine serum albumin (BSA), 1% FBS and 0.065% sodium azide] and incubated with antibodies against surface markers: CD3-APC, CD3-FITC, CD3-PB, CD16-PE/Cy7, CD4-PC5.5, CD8-APC/Cy7, CD25-PE/Cy7, CD69-APC/Cy7, CD56-PE, CD57-FITC, CD158-PE/Cy7, CXCR3-APC HLADR-APC/Cy7, NKp30-APC, NKp46-PE, NKG2D-PE, LAMP1 (CD107a)-APC, PD1 (CD279)-APC (Biolegend); CD56-PC5 and NKG2A-APC (Beckman Coulter); and, NKG2C-PE (R&D). For extracellular staining, cells were directly incubated with the appropriate conjugated antibodies at 4°C for 30 min in the dark. For intracellular staining, after surface labelling, cells were fixed with 1% p-formaldehyde for 10 min at RT, permeabilized with 0.1% saponin for 10 min at RT. After staining, cells were washed in PBA and analysed using either Gallios or CytoFLEX flow cytometers (Beckman Coulter). Analysis of the experiments was performed using Kaluza software. Statistical analyses were performed using the GraphPad Pad Prism 9 software.

### scRNA-seq

Methods followed for the sample preparation, library generation, sequencing and data analysis have been described (37). Briefly, library pools from BCG-stimulated PBMC were sequenced at 650 pM in paired-end reads on a P3 flow cell using NextSeq 2000 (Illumina) at the Genomics Unit of the National Centre for Cardiovascular Research (CNIC, Madrid, Spain). Seurat (v4.0.2) R package was used to subset and merge NK cells from the in-house BCG-primed experiment and the dataset of resting NK cells coming from (38). For visualization purposes, ViolinPlots and DoHeatmap functions from Seurat R package were used, as well as stacked_violin plots from Scanpy (v1.9.1) Python toolkit. GOBP enrichment analysis was carried out with Metascape platform (https://metascape.org/) and represented using the ggplot2 (v3.3.6) R package.

R code related to the main scRNA-seq figures can be found at GitHub (https://github.com/algarji/Felgueres_NK_scRNA-seq). scRNA-seq data from BCG-priming experiments are available at Gene Expression Omnibus (GEO): GSE203098. scRNA-seq data from resting NK cells in PBMC (38) are available at GEO: GSE149689.

### Cytokine-mediated stimulation

Aliquots of rhIL12, 15 and 21 (Peprotech) and rhIL18 (MBK) cytokines were prepared as indicated by manufacturer and stored at -80°C, so that vials were only thawed once. For minimal-dose cytokine titration experiments, PBMC cultures were stimulated by adding different concentrations of cytokines either individually or in combination. 7 days later, cells were recovered for characterization and functional assays. Different wells were recovered every seven days after cytokine stimulation.

For persistence and phenotype experiments, after a week of co-culture of PBMC and BCG, cells were re-stimulated by adding IL12, 15 and 21 to a final concentration of 0.1 ng/ml, 0.5 ng/ml, and 0.5 ng/ml, respectively. In certain cases (as indicated), IL-18 was added the day before the experiment to a final concentration of 5 ng/ml. Functional assays were performed weekly, either after BCG or cytokine stimulation, and phenotype was monitored by flow cytometry.

### Proliferation assays

PBMCs were incubated with 2 μM CellTrace™ Violet stain (Molecular Probes) for 20 min at 37LJC 5% CO_2_. Complete RPMI-1640 medium (Biowest) 5% FBS (Capricorn), 5% HS (Sigma) was then added for 5 min and the cells were washed once with complete medium before plating in 24-well plates with or without minimal-dose cytokines, either individually or in combination, as indicated. After seven days in culture, cells were recovered and analysed by flow cytometry.

### Degranulation experiments

PBMC from healthy donors were used as effector cells. Cancer cell lines, pre-treated with HP1F7 antibody, which was included in the medium at a final concentration of 10 μg/ml for 30LJmin, to block MHC-I mediated NK inhibition (unless otherwise indicated), were used as target cells (K562 cells were used as positive control). 25000 effector cells (normalizing for NK cells) were incubated with 50000 target cells (1:2 E:T ratio), unless otherwise indicated, for 2h as described (39). Surface expression of LAMP1 (CD107a) was analysed by flow cytometry. Statistical analyses were performed using the GraphPad Pad Prism 9 software.

### IFNγ-release assays

For intracellular IFNγ staining, PBMC were co-cultured with target cells for 6 hours at 1:2 E:T ratio (normalizing for NK cells) at 37LJC 5% CO_2._ After 1 hour of co-incubation, brefeldin-A (Biolegend) was added to a final concentration of 5 μg/ml. 5 hours later, cells were recovered, fixed, permeabilised, and stained with IFNγ-PE (Biolegend).

### Cytotoxicity Assays

10^4^ target cells were plated in 96-well flat-bottom plates in triplicates in a final volume of 0.2 mL and let to adhere overnight. The next day cells were labelled for 1 h with medium containing 3 µM calcein-AM (Molecular Probes), washed 3 times and incubated in fresh medium for a further hour to release free dye. Target cells were then pre-treated with HP1F7 to block MHC-I, unless otherwise indicated. Cells were resuspended in complete RPMI-1640 without phenol red (Gibco) to minimisel interference. Effector cells were incubated with adherent target cells at a 5:1 E:T ratio (except for Fig 5C, E:T corresponds to whole PBMC: target), the percentage of NK cells was determined for each donor and cell numbers adjusted accordingly for 5:1 NK:target) for 3 h at 37°C and 5% CO_2_. Supernatants were recovered, after centrifugation at 1300 rpm for 5 min to pellet cells and transferred to a clean opaque plate. Calcein-AM release was determined by measuring absorbance (excitation wave 485LJnm and emission wave 535 nm), using BioNova® F5 System. Specific lysis was calculated as the ratio [(valueLJ−LJspontaneous release) / (maximumLJrelease−LJspontaneous release)]LJ×LJ100. Spontaneous release corresponds to labelled target cells without effector cells. Maximum release was determined by lysing the target cells in 0.5% Triton X-100 (ThermoScientific). In all the experiments, the spontaneous release was between 20% and 30% of the maximum release. For some experiments, to block CD94, effector cells were pre-treated for 30 min with 10 μg/ml of F(ab’)_2_ fragments [generated according to manufacturer’s instructions (Thermo Scientific™Pierce™ F(ab’)2 Micro Preparation Kit)] of the HP3B1 antibody (kind gift of Miguel López-Botet) (40). Statistical analyses were performed using the GraphPad Pad Prism 9 software.

## Results

### BCG-stimulated CD56^high^CD16^+^ NK cells degranulate against multiple types of solid tumours

To test whether BCG-stimulated anti-tumour NK cells could kill a range of solid tumours besides bladder cancer, PBMC from healthy donors were incubated with BCG for 7 days and their ability to recognise a panel of solid tumour cell lines was evaluated. As previously observed, although the percentage of total NK cells did not increase during this week in culture, the percentage of CD56^high^ cells increased significantly compared to untreated cells (Fig. 1A, B; Fig. S1A). Since these BCG-primed cells do not correspond to peripheral blood immature CD56^bright^ NK cells, but rather, are activated NK cells we refer to them as CD56^high^. Degranulation assays against a panel of cell lines representing different types of solid tumours (Fig. 1C), showed that BCG-activated NK cells were able to efficiently recognize all these cell lines, suggesting that BCG priming could potentiate NK cell activity against multiple cancers, not just bladder cancer.

**Fig. 1.**
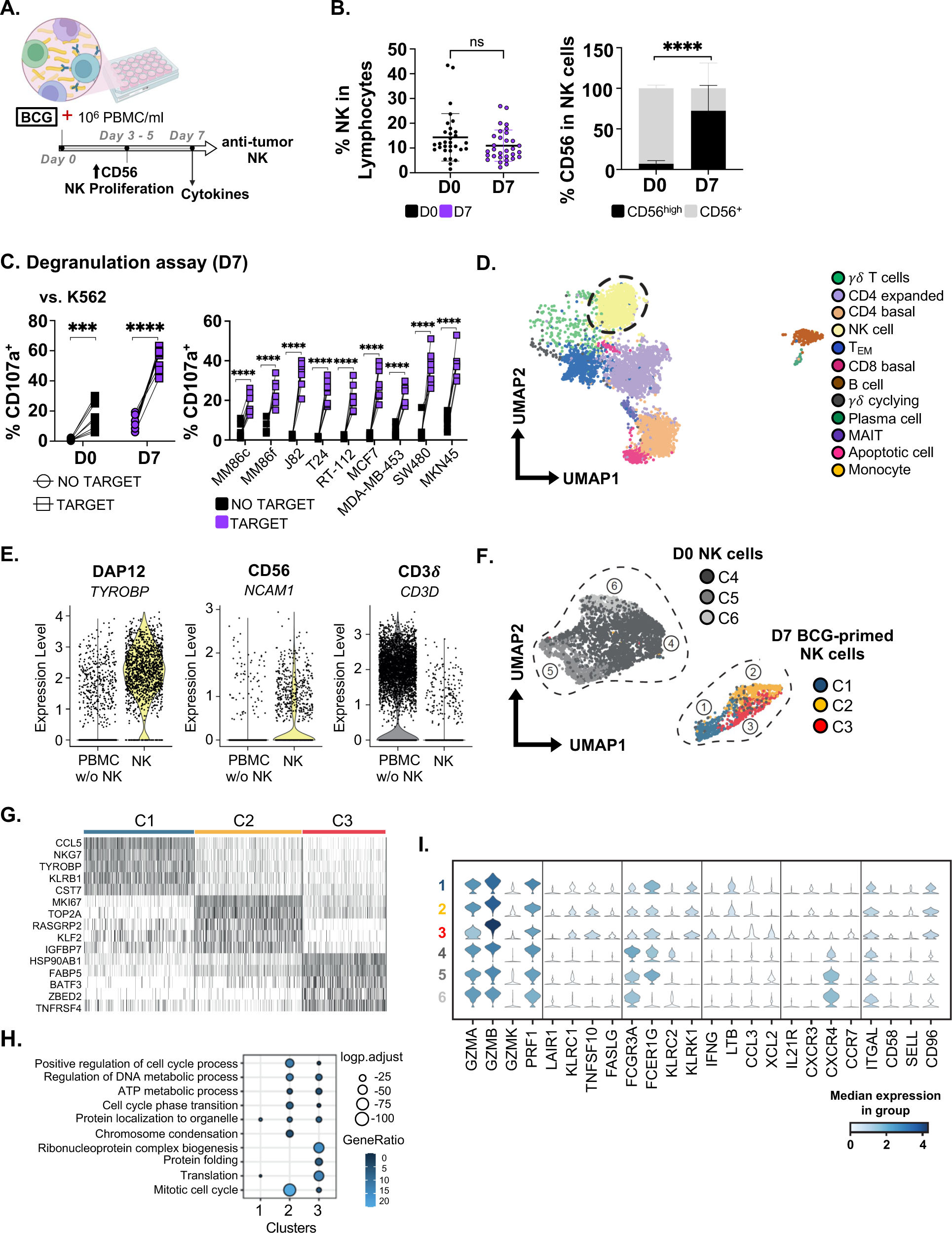
BCG-incubation leads to expansion of activated NK cells capable of killing a wide variety of solid tumour cells. **A.** Exposure of PBMC to BCG as a model for bladder cancer treatment. Previous work from the research group showed upregulation of CD56 expression marker on NK cells and increased NK proliferation *in vitro* 3-5 days after co-culture of PBMC with BCG without addition of exogenous cytokines (30). At day 7, NK cells exhibited potent degranulation against bladder cancer cells and several soluble factors were detected during the week in culture. **B. BCG-activated NK cells**. PBMC from 31 healthy donors in 15 independent experiments were co-cultured with BCG for a week. At day 7, cells were analysed by flow cytometry to determine the percentage of NK cells in the culture and the percentage of the CD56^high^ subset (black bar). Statistical analysis was done using an unpaired sample t-test (****p <0.0001). **C. Degranulation against different solid tumour cell lines**. BCG-activated PBMC were used as target (1:2 E:T ratio, NK to target) in degranulation experiments against K562 cells (positive control) or solid tumour target cell lines: melanoma (MM86c, MM86f), bladder (T24, J82, RT-112); breast (MCF7, MDA-MB-453); colon (SW480); and gastric (MKN45) cancers. Surface LAMP-1 (CD107a) was measured by flow cytometry. Statistical analysis (n=8) was done by unpaired sample t-test (****p <0.0001). D-I. scRNA-seq analysis of BCG-activated NK cells. **D. PBMC clusters**. UMAP plots represent the 12 clusters identified in BCG-activated PBMC from 3 healthy donors and analysed by scRNA-seq. A dashed circle highlights the NK cell cluster. **E. NK cluster**. Violin plots confirm the expression of NK-associated genes, *TYROBP* (DAP12), *NCAM1* (CD56), and *CD3D* (CD3δ) in the PBMC cluster (excluding NK cells) compared to NK cells. **F. Differential transcriptome of peripheral blood NK cells and BCG-primed NK cells**. UMAP represents three subclusters identified within BCG-primed NK cells (C1, C2, C3) and three subclusters identified within peripheral blood NK cells from 4 healthy donors (C4, C5, C6) (38). Subcluster annotation was performed using FindClusters and FindMarkers (see scRNA-seq in Materials and Methods). **G. Heat-map**. Heat-map represents the scaled expression of the 5 most differentially expressed genes in the three BCG-primed NK cell subclusters (C1, C2, C3). **H. Selected GO terms**. Comparison among the BCG-primed NK cell subclusters (C1, C2, C3) using markers differentially expressed within each cluster. **I. Canonical NK cell markers in BCG-activated vs resting NK cells**. Violin plots analysing the expression of NK-related molecules in BCG-primed NK cells subclusters (C1, C2, C3) compared to peripheral blood NK cell subclusters (C4, C5, C6).

In-depth characterization of BCG-primed cells from three healthy donors by scRNA-seq analysis (37) defined 12 clusters based on differential gene expression (Fig. S1B) showing, consistent with flow cytometry (30), that several populations were activated, including NK cells, CD4 T, T_EM_ and MAIT cells (Fig. 1D). The markers used to define the NK cell subpopulation in this analysis of scRNA-seq data are shown in Fig. 1E. To better define the effect of BCG priming on NK cells, these data were compared with publicly available scRNA-seq data from resting peripheral blood NK cells (38). Quality control and cell numbers of both BCG-activated and resting NK cells are shown in Fig. S1C, D, respectively. After clustering, the merged Uniform Manifold Approximation and Projection (UMAP) plots clearly showed that BCG-stimulated (C1-3) and resting (C4-6) NK cells were located separately within the 2D projection, due to their different transcriptomic signatures (Fig. 1F, Fig. S1E). Within the BCG-activated NK cells, C1 had features consistent with a non-proliferating CD56^dim^ subset while clusters C2 and C3 corresponded to highly proliferative and activated cells (Fig. 1G, H). Stacked violin plots were built to compare differential expression of key NK molecules (Fig. 1I). Cells in the C3 cluster differentially expressed genes associated with strong cytotoxic capacity, such as *GZMB* and *XCL*2. Transcripts for CD94 (*KLRC1*) and CD16 (*FCGRIIIA*) proteins identified in BCG-activated NK cells by flow cytometry were also present, although the relative abundance of RNA and protein differed somewhat.

All three clusters of BCG-primed NK cells showed enriched expression of cytotoxicity-related transcripts, such as granzymes, perforins, TRAIL (*TNFSF10*), *FASLG, IFNG, LTB, CCL3* and NKG2D (*KLRK1*). Interestingly, expression of chemokine receptor CXCR3 transcripts correlates with high concentrations of CXCL10 found in urine from BCG-treated bladder cancer patients (32) suggesting a role for the CXCL10/CXCR3 axis in the context of this therapy.

BCG-primed NK cells did not have a clearly defined memory-like phenotype, since NKG2C (*KLRC2*) was not expressed and *FCER1G* was found at levels comparable to resting cells. Overall, the data from this analysis revealed that BCG-primed NK cells show a phenotype of activation and maturity coupled with a migration capacity well suited for anti-tumour activity.

### Minimal concentrations of cytokines support proliferation of cytotoxic NK cells

It is well known that IL12, 15, 18, and 21, have different effects on proliferation and activation of NK cells. However, these cytokines were found in extremely low or undetectable concentrations in BCG-primed PBMC cultures containing functional anti-tumour NK cells (30), as well as in bladder cancer patients treated with BCG (32). These data suggest that extremely low doses of cytokines could be sufficient for NK stimulation, perhaps with the extra benefit of not causing exhaustion of these immune cells. Therefore in the next experiments we systematically established the lowest cytokine concentrations, alone and in combination, which would support proliferation and activation of cytotoxic NK cells, without exhaustion. Titration experiments were performed incubating PBMC for 7 days with just one dose of individual cytokines on day 0. NK proliferation and percentage of CD56^high^CD16^+^ NK cells were evaluated (Fig. 2A, Fig. S2A). Then, the same minimal doses were tested in combination (Fig. 2B, Fig. S2B). A minimal dose of IL21, together with IL15 and IL12 [0.1 ng/ml IL12, 0.5 ng/ml IL15, and 0.5 ng/ml IL21], yielded the highest number and proportion of NK cells, with a CD56^high^CD16^+^ phenotype in the week-long cultures. As IL18 is usually important for NK function and proliferation (13), the combination IL12, IL15, IL18 [5 ng/ml] was also tried in functional assays using a panel of solid tumour cell lines as targets (Fig. 2C, D, E). NK cells activated with either cytokine combination responded efficiently against melanoma, bladder, and breast cancer cell lines. Although both cytokine combinations led to similar phenotypes regarding CD56^high^ CD16^+^ NK cells, specific lysis capacity appeared to be overall more efficient after stimulation with IL12, 15, and 21 rather than the combination with IL18 (Fig. 2E). These data demonstrate that human NK cells can survive and remain activated for anti-tumour responses after culture in minimal doses of cytokines without survival re-stimulation for one week.

**Fig. 2:**
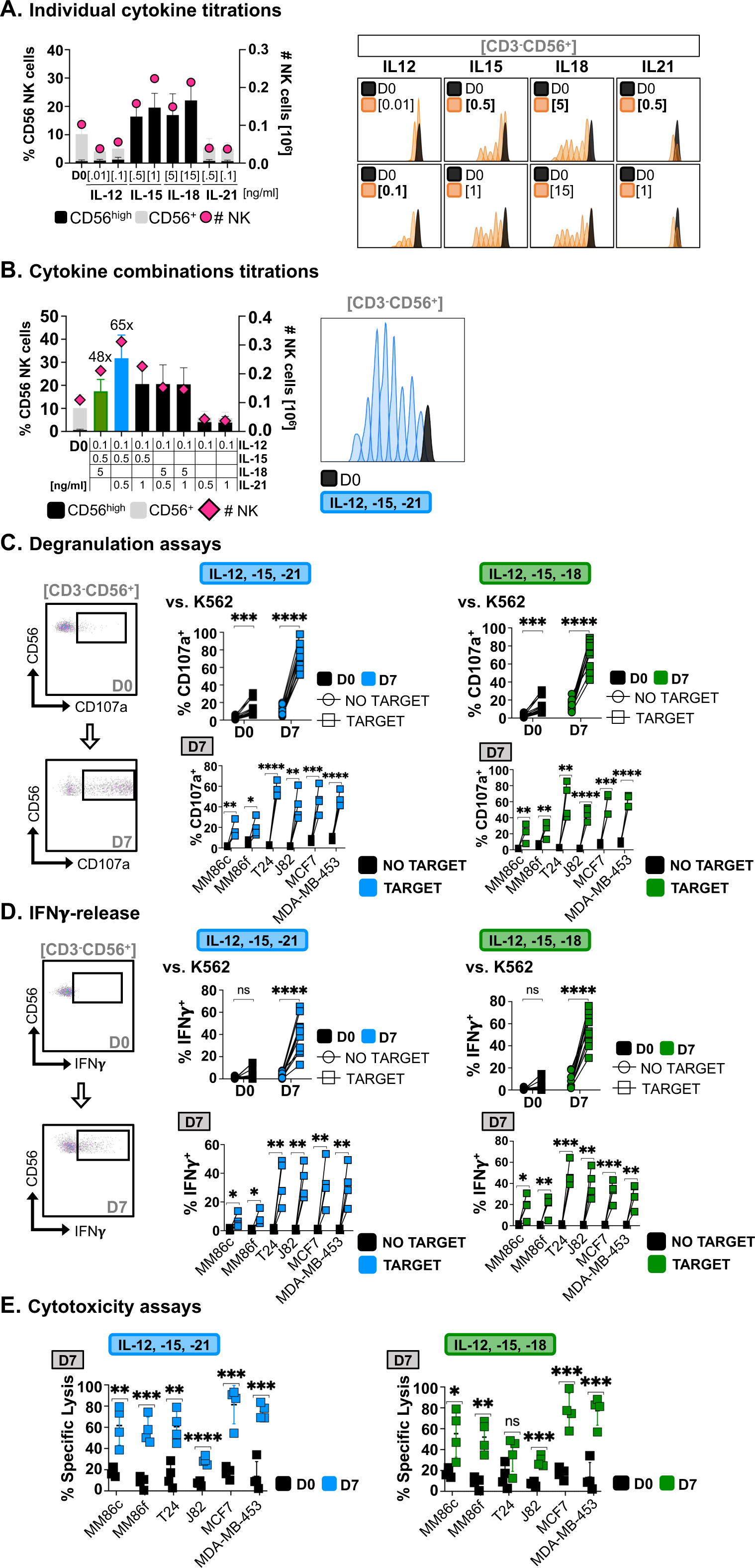
Minimal-dose IL12, 15, and 21 enhance proliferation and function of anti-tumour NK cells *in vitro*. **A. Effect of individual low-dose cytokines**. PBMC from 10 healthy donors were incubated as indicated. After a week in culture, cells were counted (pink circles, right Y axis) and the percentage of NK cells (left Y axis) was analysed by flow cytometry. Black corresponds to CD56^high^, grey to CD56^lo^. Average and standard deviation are shown. PBMC were stained with 2 µM CellTrace^TM^ Violet and NK proliferation was analysed by flow cytometry after 7 days in culture. A representative donor is shown. **B. Minimal-dose cytokine synergistic effect.** After a week in culture, BCG-stimulated cells from 10 healthy donors were counted (pink rhombus, right Y axis) and the percentage of NK cells (left Y axis) was analysed by flow cytometry. Average and standard deviation are shown. A representative NK cell proliferation plot after one-week incubation with minimal-dose IL12, 15 and 21 is shown. **C, D, E. Degranulation, IFNγ release, and cytotoxicity assays.** After cytokine activation (with the indicated combinations), NK cells from 4 healthy donors were tested as effector cells (1:2 E:T ratio, NK to target) against solid tumour target cell lines: melanoma (MM86c, MM86f), bladder (T24, J82); and breast (MCF7, MDA-MB-453) cancers. K562 cells were used as positive control (n=12). Surface LAMP-1 (CD107a) (C) and intracellular IFNγ (D) were measured by flow cytometry. Results against solid tumour targets were obtained in 2 independent experiments. Representative dot plots of degranulation and IFNγ release are shown. For cytotoxicity assays (E), effector cells were incubated with solid tumour target cells labelled with calcein-AM (5:1 E:T ratio). Dye-release was measured in 3-hour experiments and specific lysis was calculated as the ratio [(valueLJ−LJspontaneous release) / (maximumLJrelease−LJspontaneous release)]LJ×LJ100. Statistical analyses were done by unpaired sample t-tests (*p <0.05, **p <0.01, ***p <0.001, ****p <0.0001).

### BCG-priming of a PBMC culture followed by minimal-dose cytokines preferentially expands NK cells with anti-tumour capacity

Next, we investigated whether these minimal doses of cytokines could maintain NK cells in long-term cultures and how BCG-priming contributed to their anti-tumour capacity. PBMC were cultured with either a combination of minimal-dose IL12, 15, and 21, added just once a week for three weeks, or primed with BCG and, after one week in culture, stimulated with minimal-dose IL12, 15, and 21 weekly, for 2 additional weeks. Cultures stimulated only with cytokines maintained cell numbers; in contrast, cytokine addition to BCG-primed cultures doubled cell numbers (Fig. 3A, right axes), and markedly increased the percentage of activated NK cells (Fig. 3A, left axes) which was even greater after another week in culture. These cells degranulated efficiently on exposure to K562 cells, while IFNγ production was unaffected (Fig. 3B).

**Fig. 3:**
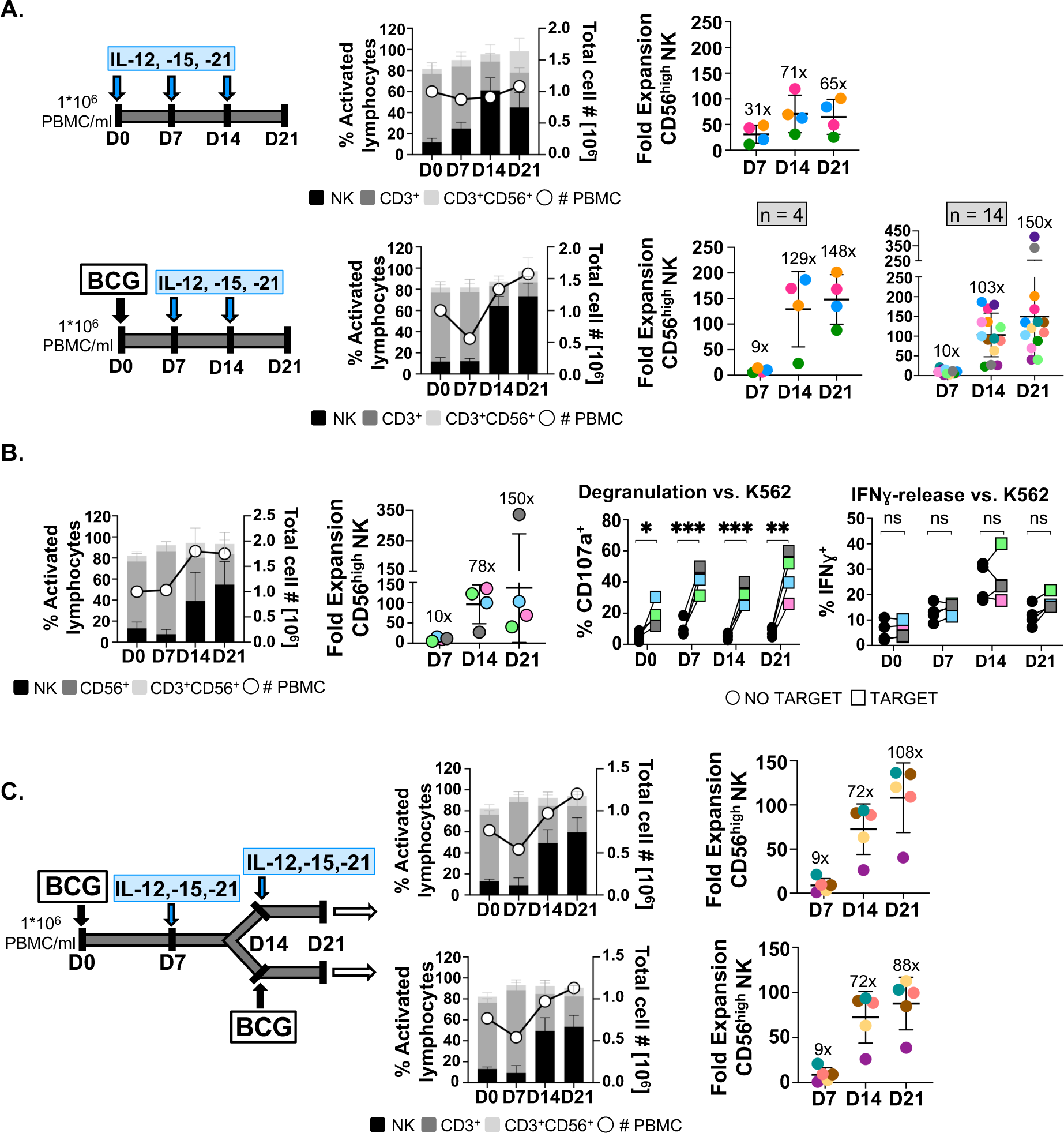
BCG-activation of PBMC followed by weekly stimulation using minimal doses of cytokines expands, after three weeks, a prominent population of cytotoxic NK cells *in vitro*. **A. Minimal-dose cytokine-primed vs**. BCG-primed NK cell expansion for 21 days. PBMC from 4 healthy donors were incubated for a week either with minimal-dose IL12, 15, and 21 or with BCG and then, weekly re-stimulated with minimal-dose cytokines between one-week resting periods (experimental design is depicted). Cells were counted (white circles, right Y axis) and analysed by flow cytometry. The percentage and standard deviation of percentage of activated lymphocytes (left Y axis) are shown as well as the different lymphocyte subsets, depicted with different shades, as indicated. CD56^high^ NK cell expansion is plotted for each donor with different colours and the average number is shown above the graph. For comparison of the two conditions, 4 donors were used. A further 10 donors were tested with the BCG followed by cytokine combination in 5 independent experiments. Expansion of effector NK cells was calculated considering the percentage of NK cells and the number of live cells in the culture; this number was compared to the initial CD56^bright^ NK cell percentage. **B. BCG-primed minimal-dose cytokine-stimulated (D21) NK cells functional assays.** PBMC from 4 healthy donors, depicted in different colours, were treated with BCG followed by weekly stimulation with minimal-dose IL12, 15, and 21 for three weeks. Cells were counted (white circles, right Y axis) and the different lymphocyte subpopulations determined by flow cytometry (left Y axis, left panel) and the effector NK cell-fold expansion was calculated (middle panel). 10^5^ NK cells were recovered weekly and tested as effector cells against K562 target cells (1:2 E:T ratio) measuring surface LAMP-1 (CD107a) and intracellular IFNγ by flow cytometry. Statistical analysis was done by unpaired t-test (*p <0.05, **p <0.01, p <0.05; ***p <0.001, ****p <0.0001). **C. BCG re-stimulation *vs.* minimal-dose cytokine re-stimulation**. PBMC from 5 healthy donors were incubated with BCG and stimulated with either minimal-dose IL12, 15, and 21 or with BCG at day 14 (see experimental design). Cells were counted (white circles, right Y axis), the different lymphocyte subpopulations were determined by flow cytometry (left Y axis), and the CD56^high^ NK cells fold expansion was calculated and depicted in different colour for each donor.

To assess whether additional BCG could improve NK cell number and stimulation, that is, whether any memory feature would enhance the response, experiments re-stimulating cultures either with BCG or cytokines one week later (day 14) were performed (Fig. 3C), showing no benefit for BCG re-stimulation. Altogether, these data demonstrate that combination of BCG priming of PBMC with minimal doses of IL12, 15, and 21 drives efficient activation and expansion of NK cells with high effector function against a range of tumour cells.

### Effector NK cells expanded after BCG-priming with minimal cytokine stimulation kill solid tumour cells after a month in culture

To further characterise the long-term persistence, phenotype and function of NK cells primed with BCG followed by minimal-dose cytokine-stimulation, PBMC from 6 healthy donors were kept in culture for one month. Cell numbers were still expanding at day 28 and NK cells were, on average, 62% of the culture (Fig. 4A), signifying an average 209-fold expansion (35- to 517-fold) from the initial number in PBMC. Each week, an aliquot of cells was recovered and used as effector cells in degranulation and cytotoxicity assays against bladder, melanoma, breast, gastric, colon and lung cancer cell lines (Fig. 4B, C). BCG-primed CD56^high^ NK cells efficiently recognized and killed a wide range of target solid tumour cells with variable intensities confirming that they kept their anti-tumour capacity even over a prolonged period of culture *in vitro*. Although tumour recognition and killing intensity varied among solid tumour cell lines and donors, these results confirm that BCG-priming followed by once-a-week minimal-dose cytokines expands activated NK cells for at least one month. These non-exhausted cultures could provide a new tool for anti-tumour effector cell expansion.

**Fig. 4:**
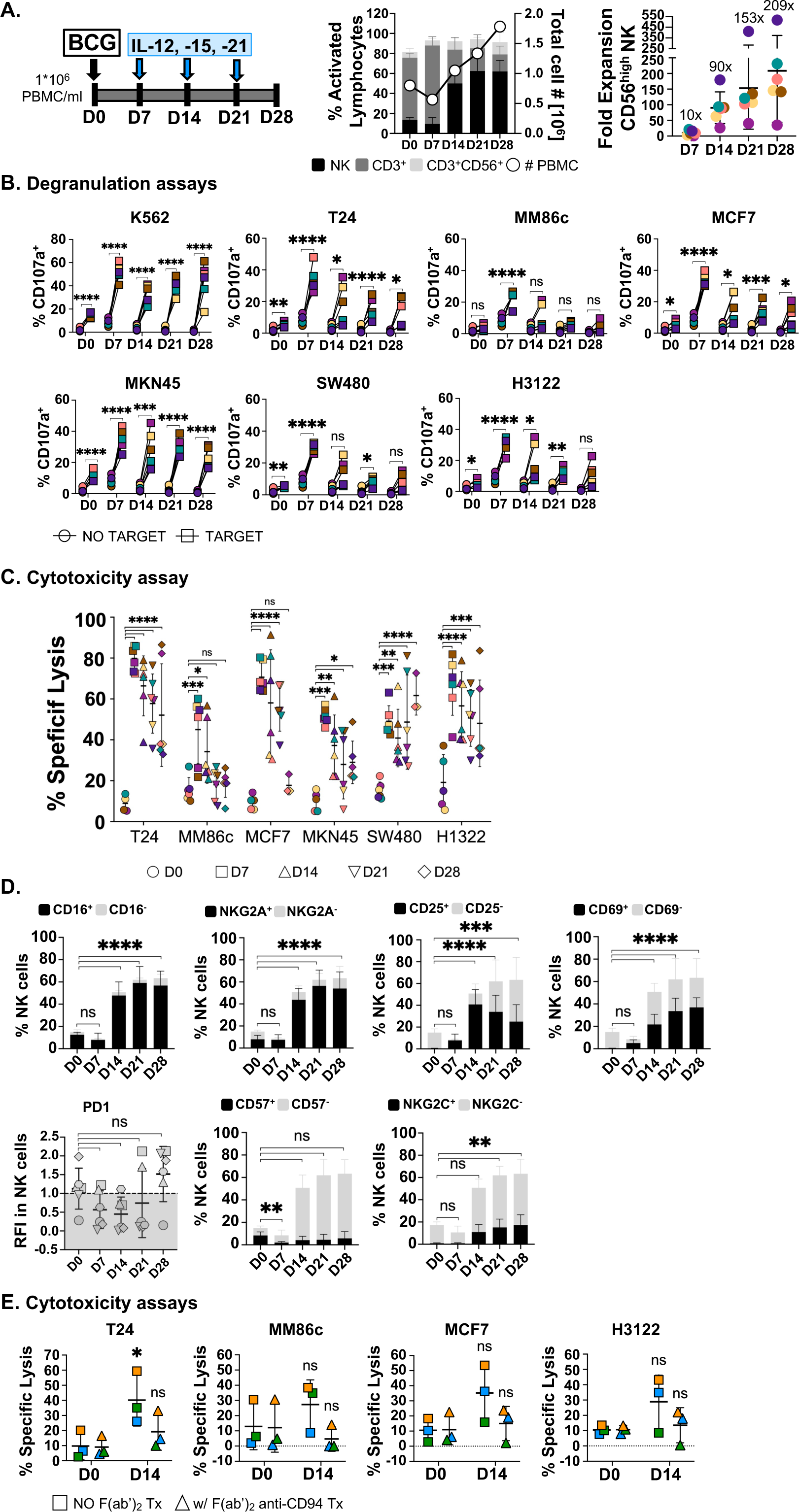
BCG-primed minimal-dose cytokine-stimulated effector NK cells proliferate for at least one month and maintain effector functions. **A. NK cell expansion after four weeks in culture.** PBMC from 6 healthy donors were incubated with BCG and stimulated with weekly minimal-dose IL12, 15, and 21 after weekly resting periods for 28 days (see experimental design). Cells were counted (white circles, right Y axis) and the different subsets analysed by flow cytometry (left Y axis). CD56^high^ NK cell expansion was calculated considering the percentage of NK cells and the number of live cells in the culture; this number was compared to the initial CD56^bright^ NK cell percentage. **B, C. Degranulation and cytotoxicity assays.** NK cells were tested as effector cells against solid tumour target cell lines bladder (T24) melanoma (MM86c), breast (MCF7), gastric (MKN45), colon (SW480), and lung (H3122) cancers. K562 were used as positive control. For degranulation (B), a 1:2 E:T ratio (NK to target) was used and surface LAMP-1 (CD107a) was measured by flow cytometry. For cytotoxicity assays (C), effector cells were incubated with solid tumour target cells labelled with calcein-AM (5:1 E:T ratio). Dye-release was measured in 3-hour experiments and specific lysis was calculated as the ratio [(valueLJ−LJspontaneous release) / (maximumLJrelease−LJspontaneous release)]LJ×LJ100. Statistical analyses were done by unpaired sample t-tests (*p <0.05, **p <0.01, ***p <0.001, ****p <0.0001). **D. NK receptors.** BCG-primed cytokine-activated NK cells were phenotype for the indicated panel of receptors. Percentage of NK cells expressing each marker is represented in black. The column height corresponds to the percentage of NK cells within the lymphocyte region in each culture. Average and standard deviation are depicted. For PD1, because the whole NK population had a single peak, the different levels of expression are shown as relative fluorescence intensity (RFI): RFI = MFI sample / MFI CD3^-^CD56^-^ (negative control), where MFI is mean fluorescence intensity. An RFI>1 means above the negative control. Statistical analysis of NK cells expressing each marker against basal expression (D0) was done by unpaired t-test (*p <0.05, **p <0.01, p <0.05; ***p <0.001, ****p <0.0001). **E. Specific cytotoxicity in the presence of anti-CD94 antibody (Fab.** Effector cells from the 3 healthy donors from (D) were incubated with solid tumour target cells: bladder (T24) melanoma (MM86c), breast (MCF7), gastric (MKN45), colon (SW480), and lung (H3122) cancers. When indicated, effector cells were pre-treated with the anti-CD94 antibody HP3B1. Target cells were not pre-treated with HP1F7. Target cells were labelled with calcein-AM (5:1 E:T ratio). Dye-release was measured in 3-hour experiments and specific lysis was calculated as the ratio [(valueLJ−LJspontaneous release) / (maximumLJrelease−LJspontaneous release)]LJ×LJ100. Statistical analyses were done by unpaired sample t-tests (*p <0.05, **p <0.01, ***p <0.001, ****p <0.0001).

The phenotype of these long-lived effector NK cells was characterized by flow cytometry (Fig. 4D). As we previously described for one-week BCG-primed PBMC (30), CD56^high^ NK cells also expressed CD16^+^ and NKG2A^+^ at day 28, after maintaining the culture with minimal doses of cytokines. Activation was confirmed by expression of CD25 and CD69 by a high percentage of NK cells, although CD25 started to decrease after day 21. These long-lived effector NK cells did not show any signs of senescence, as they continued to proliferate for the whole month. At day 14, HLA-DR and the homing receptor CXCR3 were expressed in a significantly higher proportion of NK cells. Even, at day 28, PD-1 expressing cells represented at most 2% of the NK cell population. The expression of NKp46, NKp30 and NKG2D did not change significantly after two weeks (Fig. S3). Interestingly, after minimal-dose cytokine stimulation of BCG-primed NK cells, around 22% were positive for NKG2C at day 14, and this percentage was even higher the following weeks. Treatment with F(ab’)_2_-fragments of anti-CD94 decreased cytotoxicity mediated by the mixed-lymphocyte culture against a range of different tumour types (Fig. 4E), suggesting that NKG2A inhibition is not playing a crucial role in this system.

Taken together, these results confirm that exposure of PBMC to BCG followed by weekly stimulations with minimal-dose IL12, 15, and 21 cytokines dramatically expands a cytotoxic, long-lived CD56^high^CD16^+^NKG2A^+^ NK cell subset which remain fit for target recognition and killing after 21 days in culture without reaching exhaustion. All together these data suggest that BCG priming followed by minimal dose cytokine stimulation expands a population of activated NK cells with strong anti-tumour cytotoxic capacity, independently of the expression of the inhibitory receptor CD94/NKG2A (31), which could be further exploited to develop a universal allogeneic NK cell therapy against solid tumours.

### Anti-tumour NK cells can be expanded using mycobacteria cell wall fractions and cytokine combinations

The BCG preparations used in our experiments contain only around 2% live bacteria (34). Previously, however, we have shown that NK activation could also be achieved by incubation with different mycobacteria-derived extracts for one week. So, we tested activation, proliferation and function of cell cultures stimulated with either BCG or the cell wall fraction of *M. bovis* and further expanded in minimal doses of cytokines (Fig. 5). Although cell numbers in both cultures were comparable at day 14, BCG-primed cultures contained 60-70% NK cells, while cell wall-stimulated cultures had around 40% NK cells in average. However, the phenotype of NK cells activated in both conditions, was similar with expression of CD16, CD25 and NKG2A (Fig S3B). Interestingly, cell wall stimulated NK cells degranulated very efficiently against K562 cells and solid tumours, even more strongly than BCG-primed cells for some tumour targets.

**Fig. 5:**
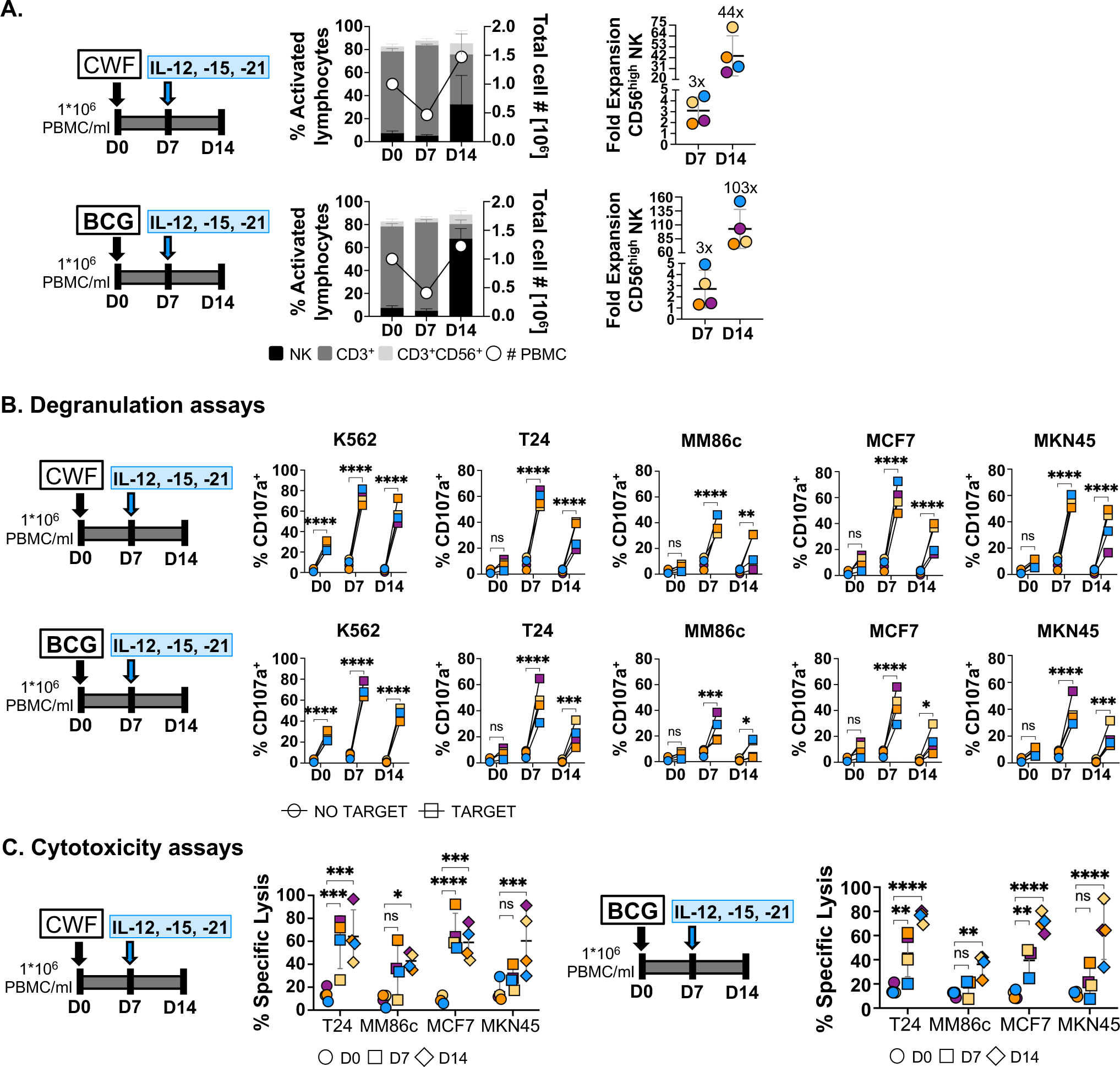
NK cells proliferate after priming with cell wall fractions followed by minimal-dose cytokine-stimulation and acquire enhanced effector functions. **A. NK cell expansion after two weeks in culture.** PBMC from 4 healthy donors were incubated with either BCG or with cell wall fraction of *M. bovis* (CWF) and stimulated after one week resting period with minimal-dose IL12, 15, and 21 (see experimental design). Cells were counted (white circles, right Y axis) and the different subsets analysed by flow cytometry (left Y axis). CD56^high^ NK cell expansion was calculated considering the percentage of NK cells and the number of live cells in the culture; this number was compared to the initial CD56^bright^ NK cell percentage. **B, C. Degranulation and cytotoxicity assays.** NK cells were tested as effector cells against solid tumour target cell lines bladder (T24) melanoma (MM86c), breast (MCF7) and gastric (MKN45) cancers. K562 were used as positive control. For degranulation (B), a 1:2 E:T ratio (NK to target) was used and surface LAMP-1 (CD107a) was measured by flow cytometry. For cytotoxicity assays (C), PBMC were incubated with solid tumour target cells labelled with calcein-AM (30:1 E:T ratio). Dye-release was measured in 3-hour experiments and specific lysis was calculated as the ratio [(valueLJ−LJspontaneous release) / (maximumLJrelease−LJspontaneous release)]LJ×LJ100. Statistical analyses were done by unpaired sample t-tests (*p <0.05, **p <0.01, ***p <0.001, ****p <0.0001).

Thus, using just mycobacterial fractions could provide an additional tool to generate anti-tumour NK cells.

## Discussion

Activated anti-tumour CD56^high^ CD16^+^ NK cells are generated after culture of PBMC with BCG for one week *in vitro* (30, 41, 42). The results of the present study support the hypothesis that mycobacteria activation followed by minimal doses of cytokines confers a specialised phenotype to NK cells, which acquire a precise competence for tumour recognition. A first key finding of this work is that, after BCG priming, weekly maintenance with minimal doses of IL12, 15 and 21 allows the growth and expansion (200x), for at least one month, of functional anti-tumour NK cells able to recognise several types of solid tumours. The combination did not include IL18 that is often used to enhance NK cell function, however, here this cytokine did not seem to be necessary. The use of extremely low concentrations of cytokines for the expansion of these NK cells was motivated by the observation of very low levels of soluble factors released from BCG-activated PBMC *in vitro* and in urine of BCG-treated bladder cancer patients (32), together with the possibility that excessive activation can, in many instances, result in cell anergy. Our characterisation of these BCG-primed effector NK cells showed that, while they share some features with CIML NK cells, the subset described here did not need cytokine re-stimulation to recognise tumours. Although some reports demonstrate that intravenous BCG is not toxic in primates, systemic use of whole bacteria could be worrying in oncologic patients. However, the use of defined mycobacteria fragments for NK cell priming opens the door to use this protocol without live bacteria. The strategy for the expansion of effector anti-tumour NK cells presented here has potential practical implications, establishing conditions to increase the number and fitness of cytotoxic NK cells against a variety of solid tumours, as desired for adoptive cell therapies.

The results presented here demonstrate that BCG-primed cytotoxic CD56^high^ CD16^+^ NKG2A^+^ NK cells remain fit for target recognition and killing over extended times in culture without becoming exhausted. This outcome was achieved by decreasing the cytokine concentration to the lowest possible allowing cell survival. NK cells in culture were stimulated just once weekly with only IL12 [0.1 ng/ml], IL15 [0.5 ng/ml] and IL21 [0.5 ng/ml]. This represents a lowering of the concentration at least 100 times comparing to earlier protocols of NK cell activation (17) and around 10-100 times lower than in the generation of CIML NK cells (19) and replaces IL18 with IL21. Interestingly, when briefly stimulating the BCG-primed, minimal-dose cytokine-stimulated CD56^high^ NK cells with 5 ng/ml IL18, no significant difference in either tumour cell recognition or specific lysis capacity was observed (Fig. S4). Importantly, compared to CIML, no IL15 survival boost was added at intermediate times, which would suggest good *in vivo* persistence of these cells and avoid the cell dysfunction associated with chronic cytokine exposure (43).

The minimal-dose cytokine combination has several advantages, when compared to other cytokine-mediated stimulation strategies for *ex vivo* expansion of NK cells. First, cultures start with PBMC and, without any cell sorting, resulted in 80% NK cell cultures, simplifying enormously handling time and procedures. Second, minimal use of each cytokine, together with the weekly frequency provides an important cost-related advantage. Further, the method enhanced cytotoxic function and proliferation rates while maintaining cells in a healthy, non-exhausted state, demonstrated by expression of CD25 and CD69, and no increase of PD-1 could be detected over time. This finding points to long-term survival even in resting environments, a very desirable for *in vivo* translation. Third, this strategy does not need transfection of a cytokine membrane-bound construct for expansion of effector NK cells with anti-tumour phenotype. It is also noteworthy that BCG-primed NK cells could efficiently kill multiple different types of tumours including melanoma, breast, gastric and colon cancer as shown in both degranulation and cytotoxicity assays.

The phenotype of cytotoxic BCG-primed NK cells is interesting; most of these cells express NKG2A and CD16, consistent with previous results demonstrating the anti-tumour activity of NKG2A^+^ NK cells (19, 31). It is intriguing that anti-tumour activated NK cells express NKG2A which associates with CD94 to form heterodimers with inhibitory function, upon recognition of HLA-E/peptide complexes (44, 45). Although NKG2A has been associated with a lower capacity of NK cells to recognize tumour cells, due to its inhibitory function leading to lower cytotoxic function and IFNγ production (46, 47, 48, 49), BCG-primed NKG2A^+^ NK cells are clearly still functional, like other examples of NKG2A^+^ NK cells described in the literature (31, 50). Interestingly, increased expression of NKG2A was also observed in the generation of CIML NK cells, where enhanced memory-like IFNγ production was associated with less mature NK cells that responded better to cytokine-mediated stimulation (19). Here, functional data demonstrate that the NK subset generated after BCG-priming acquired a cytotoxic anti-tumour phenotype which decreased upon CD94 blockade, suggesting that NKG2A is not inhibiting these cells, but rather that the activating CD94/NKG2C, expressed by around 20% of these NK cells, could be important for tumour recognition. Detailed studies on the role of the CD94/NKG2A-C heterodimers in this context are underway.

The present study demonstrates that it is possible to generate large numbers of cytotoxic NK cells with broad anti-tumour recognition capacity, using BGC-primed PBMC. While the number of activated NK cells increased between 90 times (one week after cytokine stimulation) and 200 times (28 days after starting the culture), these conditions could be still improved in future research. The use of minimal cytokine concentrations just once a week highlights the possibility of these cell cultures to endure resting conditions, devoid of stimulation for long periods of time. Similarly, other conditions (reactivation time and cytokine regime) using cell wall preparations of *M. bovis* could improve NK number in these cultures. In future studies, it will be important to address effector NK cell fitness, target-specificity and longevity *in vivo*.

Overall, our results provide a first step towards the use of BCG and minimal doses of cytokines for the generation, activation, and maintenance of anti-tumour NK cells which could be potentially translated as a strategy for developing allogeneic cell agents as immunotherapy against solid tumours.

## List of Supplementary Figures

**Fig. S1.** Characterization of BCG-primed PBMC.

**Fig. S2.** Lymphocyte populations generated after one week cultured with different concentrations of individual cytokines and their combinations.

**Fig. S3.** Flow cytometry. A. BCG-primed NK cells, then stimulated with minimal-dose cytokines. B. Cell wall fraction-primed, then stimulated with minimal-dose cytokines.

**Fig. S4:** Effect of low-dose IL-18 on NK cells BCG-primed followed by minimal-dose cytokines.

## Declarations

### Ethics approval and consent to participate

Work followed the World Medical Association’s Declaration of Helsinki. PBMCs from buffy coats of healthy donors were obtained from the Regional Transfusion Centre, Madrid, with informed consent from the participants and approved by the institutional committees: Regional Transfusion Centre (PO-DIS-09) and assessed by the bioethics committee of CSIC.

### Consent for publication

Consent was obtained by the transfusion centre.

### Availability of data and material

Data were generated by the authors and available on request.

R code related to the main scRNA-seq figures can be found at GitHub (https://github.com/algarji/Felgueres_NK_scRNA-seq). scRNA-seq data from BCG-priming experiments are available at Gene Expression Omnibus (GEO): GSE203098. scRNA-seq data from resting NK cells in PBMC (38) are available at GEO: GSE149689

### Competing interests

The authors declare no competing interests.

### Funding

Spanish Ministry of Science and Innovation [Ministerio de Ciencia, Innovación y Universidades (MCIU)/Agencia Estatal de Investigación (AEI)/European Regional Development Fund (FEDER, EU)]: PID2021-123795OB-I00 (MVG), PID2020-115506RB-I00 (HTR)]; INPhINIT Doctoral Programme from La Caixa Foundation LCF/BQ/DI19/11730039 (MJF). Spanish Ministry of Science and Education fellowship FPU18/01698 (AFGJ).

### Authors’ contributions

MJF, GE, AFGJ, HTR acquired, analysed, and interpreted data. HTR, AD, LMP and MVG provided material support. MJF and MVG wrote the manuscript with critical revisions by all authors. MVG supervised the study.

## Acknowledgements

The authors would like to thank Miguel López-Botet (IMIM, Barcelona) for sharing antibody and for a critical review of the manuscript; Prof. Annette Paschen (University Hospital of Essen, Germany) for human melanoma cell lines; and Alberto Benguría and Enrique Vázquez at the Genomics Unit of the National Centre for Cardiovascular Research (CNIC, Madrid, Spain) for their advice on scRNAseq analysis; BEI Resources, NIAID, NIH for *Mycobacterium bovis*, Strain AF 2122/97 (ATCC ® BAA-935™), Cell Wall Fraction, NR-31213.

The group of MVG belongs to the research network cancer hub-CSIC. MJF and AFGJ are registered PhD students at the Molecular Biosciences doctoral program of the Universidad Autónoma de Madrid (UAM).

## List of abbreviations

ADCC: Antibody-dependent cellular cytotoxicity
BCG: Bacillus Calmette-Guérin
BSA: Bovine serum albumin
CIML: Cytokine induced memory-like
DMSO: Dimethyl sulfoxide
FBS: Fetal Bovine Serum
FDA: Food & Drug Administration
GVHD: Graft-versus-host disease
HCMV: Human cytomegalovirus HS Human serum
IFNγ: Interferon γ
IL: Interleukin
MAIT: Mucosal-associated invariant T cells
NK: Natural killer
NMIBC: Non-muscle invasive bladder cancer
PBMC: Peripheral blood mononuclear cells
PBS: Phosphate buffered saline
rhIL: Recombinant human interleukin
scRNA-seq: Single cell RNA sequencing
UMAP: Uniform manifold approximation and projection

## Notes

https://github.com/algarji/Felgueres_NK_scRNA-seq

### Competing Interest Statement

The authors have declared no competing interest.

