## Supplemental Figures and legends for "Long-term cytotoxic NK cells with broad anti-tumour capacity proliferate selectively, without exhaustion, after BCG priming and extremely low doses of cytokines": 230518 Supplementary Figure legends.docx

**Fig. S1. Characterization of BCG-primed PBMC.**

**A. Gating strategy for the selection of lymphocyte subpopulations.** Freshly thawed PBCM were analysed by flow cytometry. NK cells, CD3^+^ lymphocytes and double positive cells were selected, as indicated, within the lymphocyte region in FSC vs SSC. At day 7, BCG-activated lymphocytes were selected in the FSC vs SSC plot. Activated CD56^high^ NK cells were gated separately from the rest of CD3^−^CD56^+^ NK cells. **B. Differential gene expression in BCG-activated PBMC.** Heat-map of the whole-data set representing a scaled expression of the top 10 differentially expressed genes across the 12 clusters identified in BCG-activated PBMC from 3 healthy donors. **C, D.** scRNA-seq quality control of the BCG-activated and freshly isolated NK dataset. BCG-activated PBMC from 3 healthy donors (C) and PBMC NK cells from 4 healthy donors [37] (D) were analysed by scRNA-seq. Cells were filtered to retain between 200 and 5000 features, and present less than 10% of mitochondrial content. Doublets and negative cells were additionally removed based on the detection of HTO signal (middle). Tables summarize the number of events per sample and cluster. **E. Differential gene expression comparing BCG-activated and freshly isolated NK cells.** Heat-map of the whole-data set representing a scaled expression of the top 5 differentially expressed genes across the different NK clusters from BCG-primed NK cells (C1, C3, C3) and PBMC NK cells (C4, C5, C6).

**Fig. S2. Lymphocyte populations generated after one week cultured with different concentrations of individual cytokines and their combinations.**

PBMC from 10 healthy donors were incubated with minimal doses of cytokines (A) and combinations (B) as indicated. After a week in culture, cells were counted (pink circles or rhombus, right Y axis) and analysed by flow cytometry. The percentage and standard deviation of the different lymphocyte subsets are depicted with different colour in the bars, indicating the percentage of activated lymphocytes (left Y axis). The two cytokine combinations which triggered proliferation and CD56 upregulation the most are highlighted in green and blue.

**Fig. S3. Flow cytometry receptor characterization. A. BCG-primed NK cells stimulated with minimal-dose cytokines.** PBMC from 7 healthy donors (n=4 for NKG2D) were incubated with BCG for a week and then minimal-dose IL12, 15, and 21 cytokines were added. Cells were analysed by flow cytometry. Percentage of NK cells expressing each marker and standard deviation are plotted in black filling. Grey bars represent the average percentage of the NK subset in the activated lymphocyte gate. Statistical analysis of NK cells expressing each marker against basal expression (D0) was done by unpaired t-test (*p <0.05, **p <0.01, p <0.05; ***p <0.001, ****p <0.0001). Relative fluorescence intensity (RFI) was used to analyse the different levels of expression when the whole population was a single peak. An RFI>1 means above the negative control: the expression on CD3-CD56- subpopulation (RFI = MFI sample / MFI CD3-CD56-, where MFI is mean fluorescence intensity). **B. Cell wall fraction-primed NK cells stimulated with minimal-dose cytokines.** PBMC from 4 healthy donors were incubated with 5 µg/ml of cell wall fraction from of *M. bovis* (CWF). After a week in culture, minimal-dose IL12, 15, and 21 cytokines were added. Cells were analysed by flow cytometry. Percentage of NK cells expressing each marker and standard deviation are plotted in black filling. Grey bars represent the average percentage of the NK subset in the activated lymphocyte gate. Statistical analysis of NK cells expressing each marker against basal expression (D0) was done by unpaired t-test (*p <0.05, **p <0.01, p <0.05; ***p <0.001, ****p <0.0001).

**Fig. S4: Effect of low-dose IL-18 on NK cells BCG-primed followed by minimal-dose cytokines.**

**A. Experimental design.** PBMC from 6 healthy donors were incubated with BCG and stimulated with minimal-dose IL12, 15, and 21 after weekly resting periods, as indicated. The day before functional assays, cells were stimulated overnight with IL18 (5 ng/ml). **B, C. Degranulation and cytotoxicity assays.** For degranulation, 105 NK cells were tested as effector cells against solid tumour target cell lines (1:2 E:T ratio, NK to target): bladder (T24) melanoma (MM86c), breast (MCF7), gastric (MKN45), colon (SW480), and lung (H3122) cancers. K562 was used as positive control. Surface LAMP-1 (CD107a) (B) was measured by flow cytometry. Results were obtained in 2 independent experiments. For cytotoxicity assays (C), effector cells were incubated with solid tumour target cells labelled with calcein-AM (5:1 E:T ratio). Dye-release was measured in 3-hour experiments and specific lysis was calculated as percentage of spontaneous release. Specific lysis was calculated as the ratio [(value − spontaneous release) / (maximum release− spontaneous release)] × 100. Statistical analyses were done by unpaired sample t-tests (*p <0.05, **p <0.01, ***p <0.001, ****p <0.0001). Different donors are represented by different colours.
