## Supplemental Figures and legends for "Long-term cytotoxic NK cells with broad anti-tumour capacity proliferate selectively, without exhaustion, after BCG priming and extremely low doses of cytokines": 230518 Supplementary Figures.pdf

A.

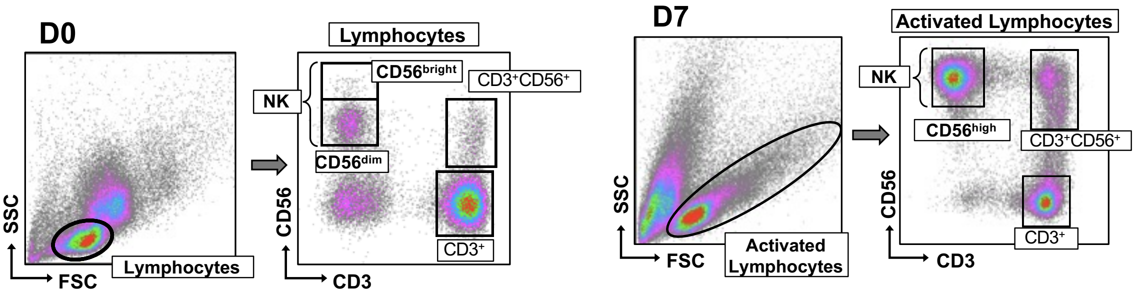


B.

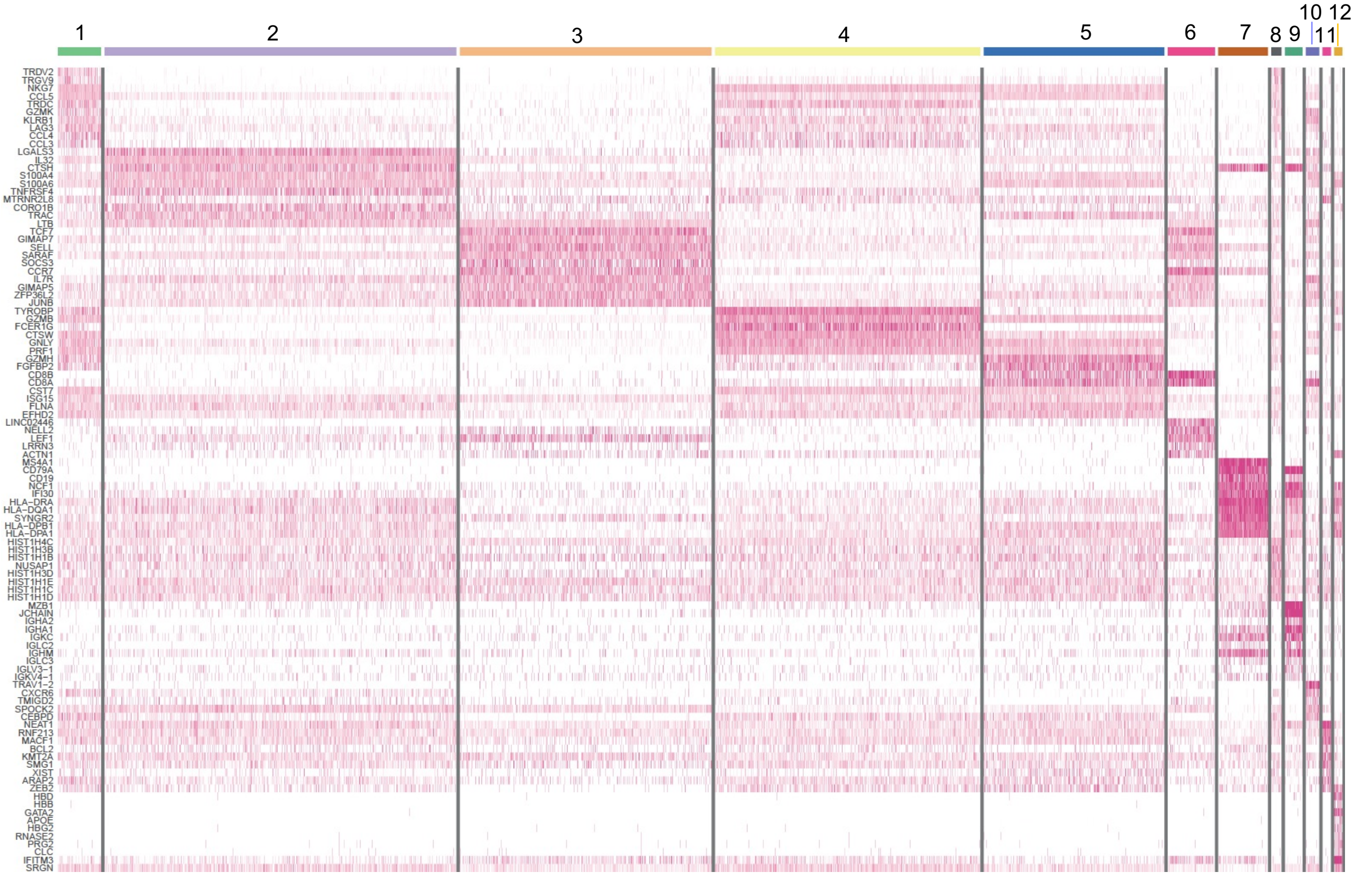


- **1:** $\gamma\delta$ T cells
- **2:** CD4 expanded
- **3:** CD4 basal
- **4:** NK cell
- **5:** $T_{EM}$
- **6:** CD8 basal
- **7:** B cell
- **8:** $\gamma\delta$ cyclying
- **9:** Plasma cell
- **10:** MAIT
- **11:** Apoptotic cell
- **12:** Monocyte

**C.**

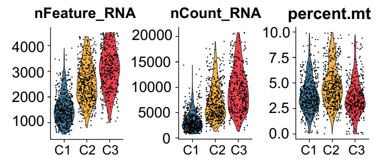


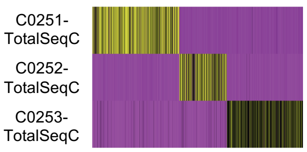


Demultiplexing

| | Sample 1 | Sample 2 | Sample 3 | TOTAL |
|---|---|---|---|---|
| **C1** | 255 | 146 | 80 | 481 |
| **C2** | 155 | 75 | 235 | 465 |
| **C3** | 131 | 74 | 157 | 362 |
| **TOTAL** | 541 | 295 | 472 | 1308 |

**D.**

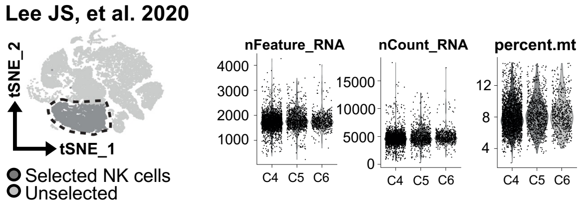


Demultiplexing

| | HD1 | HD2 | HD3 | HD4 | TOTAL |
|---|---|---|---|---|---|
| **C4** | 392 | 642 | 336 | 1088 | 1458 |
| **C5** | 296 | 244 | 223 | 106 | 870 |
| **C6** | 308 | 223 | 147 | 2 | 680 |
| **TOTAL** | 996 | 1109 | 706 | 1196 | 4007 |

**E.**

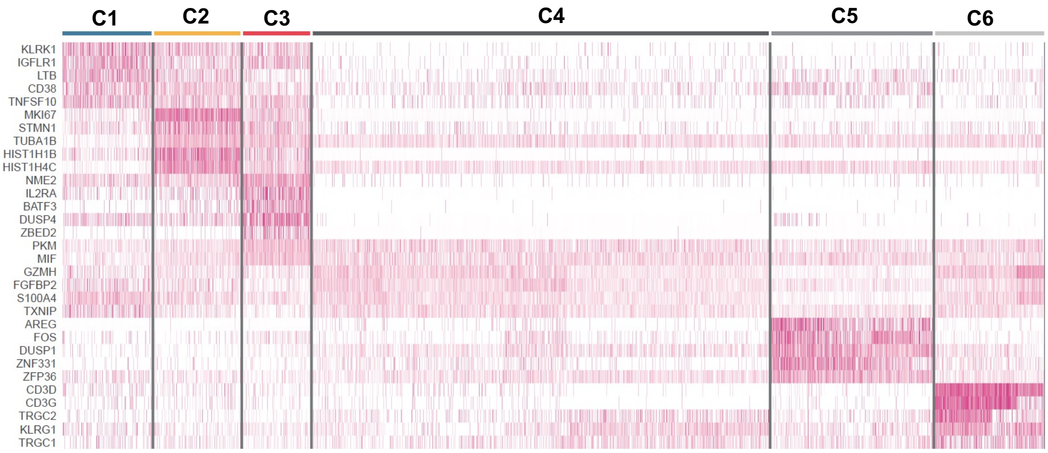

**Fig. S2**

**A.**

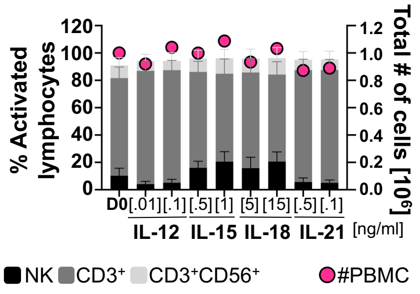

% Activated lymphocytes
Total # of cells [10^6]
120
100
80
60
40
20
0
1.2
1.0
0.8
0.6
0.4
0.2
0.0
D0 [.01] [.1] [.5] [1] [5] [15] [.5] [.1]
IL-12 IL-15 IL-18 IL-21 [ng/ml]
NK CD3+ CD3+CD56+ #PBMC


**B.**

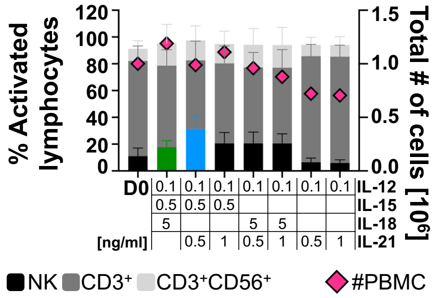

% Activated lymphocytes
Total # of cells [10^6]
120
100
80
60
40
20
0
1.5
1.0
0.5
0.0
D0
0.1 0.1 0.1 0.1 0.1 0.1 0.1 IL-12
0.5 0.5 0.5 IL-15
5 5 5 IL-18
[ng/ml] 0.5 1 0.5 1 0.5 1 IL-21
NK CD3+ CD3+CD56+ #PBMC

### Fig. S3

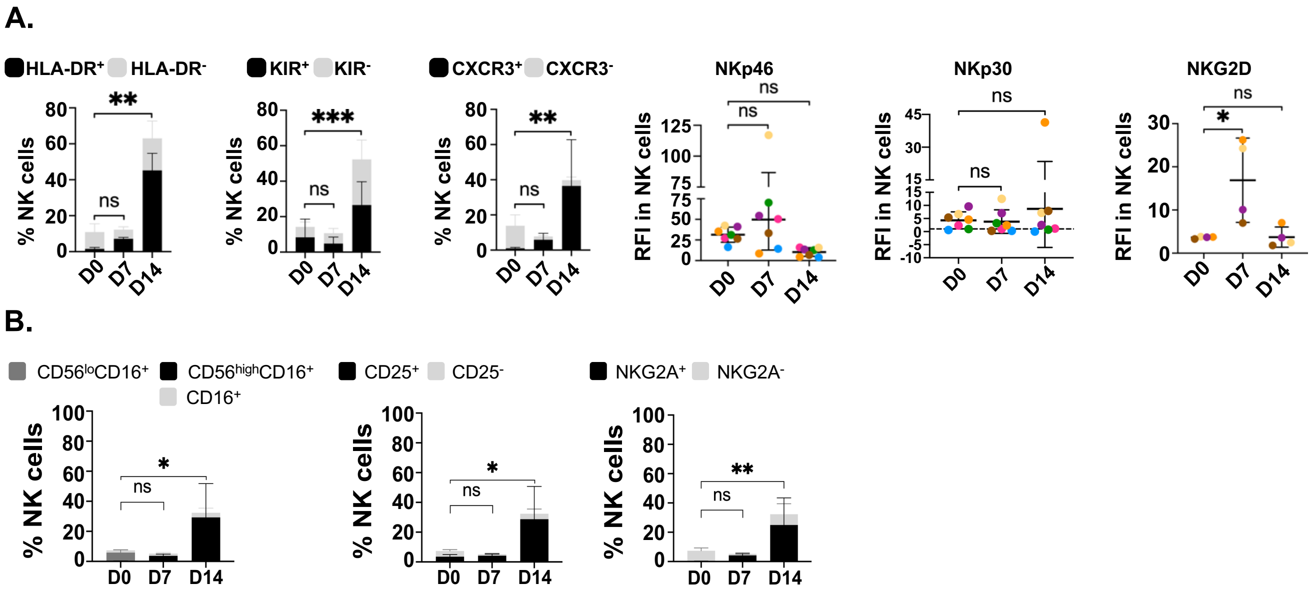
A.
HLA-DR+ HLA-DR-
KIR+ KIR-
CXCR3+ CXCR3-
NKp46
NKp30
NKG2D
% NK cells
RFI in NK cells
D0
D7
D14
ns
*
**
***
B.
CD56loCD16+
CD56highCD16+
CD16+
CD25+ CD25-
NKG2A+ NKG2A-
% NK cells
D0
D7
D14

Fig. S4

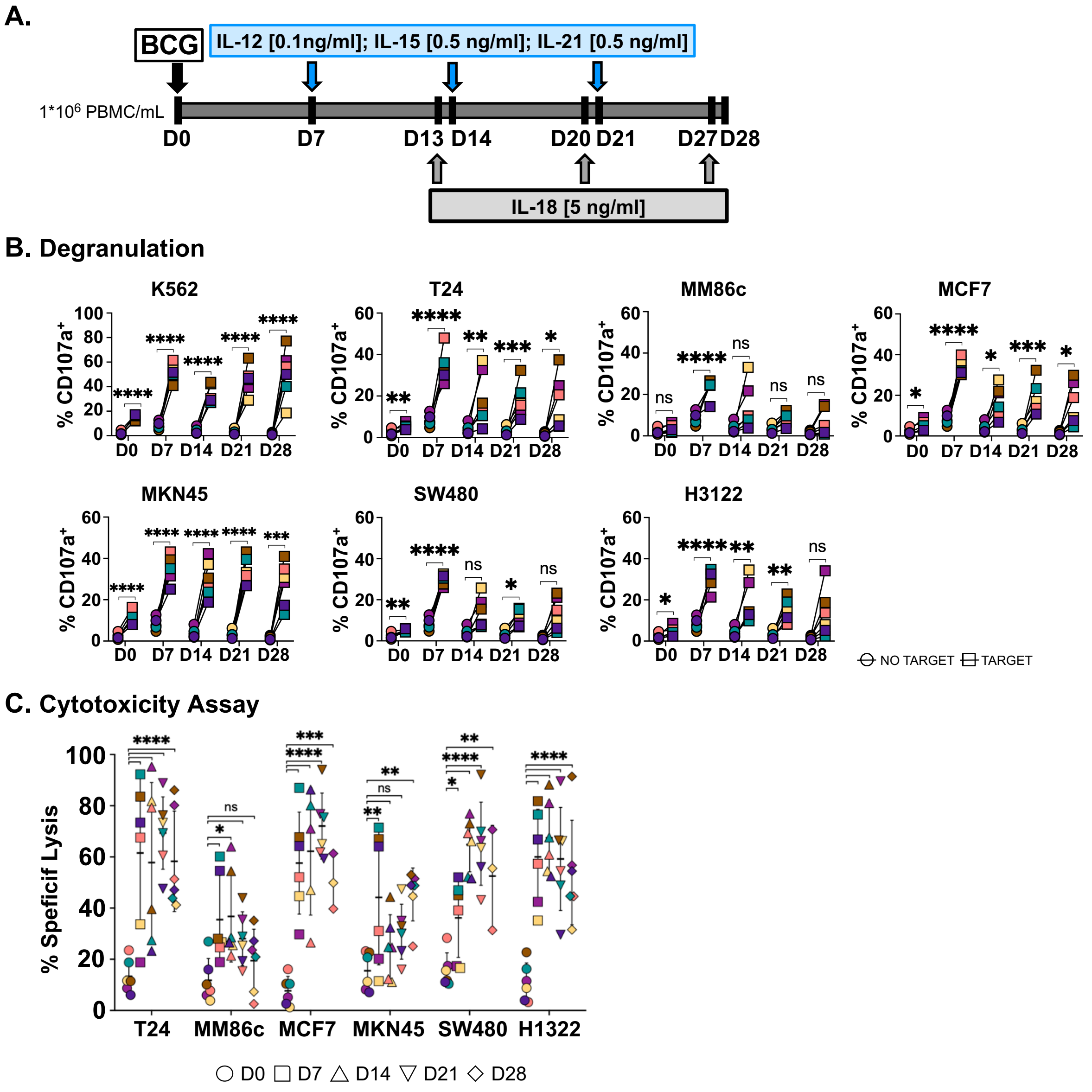
A.
BCG
IL-12 [0.1ng/ml]; IL-15 [0.5 ng/ml]; IL-21 [0.5 ng/ml]
1*10^6 PBMC/mL
D0
D7
D13 D14
D20 D21
D27 D28
IL-18 [5 ng/ml]
B. Degranulation
K562
T24
MM86c
MCF7
MKN45
SW480
H3122
% CD107a+
NO TARGET
TARGET
C. Cytotoxicity Assay
% Speficif Lysis
T24
MM86c
MCF7
MKN45
SW480
H1322
D0 D7 D14 D21 D28
